# MIReVTD, a Minimum Information Standard for Reporting Vector Trait Data

**DOI:** 10.1101/2025.01.27.634769

**Authors:** Sadie J. Ryan, Paul J. Huxley, Catherine A. Lippi, Samraat Pawar, Lauren Cator, Samuel S.C. Rund, Leah R. Johnson

## Abstract

Vector-borne diseases pose a persistent and increasing challenge to human, animal, and agricultural systems globally. Mathematical modeling frameworks incorporating vector trait responses are powerful tools to assess risk and predict vector-borne disease impacts. Developing these frameworks and the reliability of their predictions hinge on the availability of experimentally derived vector trait data for model parameterization and inference of the biological mechanisms underpinning transmission. Trait experiments have generated data for many known and potential vector species, but the terminology used across studies is inconsistent, and accompanying publications may share data with insufficient detail for reuse or synthesis. The lack of data standardization can lead to information loss and prohibits analytical comprehensiveness. Here, we present MIReVTD, a Minimum Information standard for Reporting Vector Trait Data. Our reporting checklist balances completeness and labor-intensiveness with the goal of making these important experimental data easier to find and reuse, without onerous effort for scientists generating the data. To illustrate the standard, we provide an example reproducing results from an *Aedes aegypti* mosquito study.

## Introduction

Biological data are increasing in size and scope, and the means of reporting experimental or measured data are wide ranging in format - from journals ^1^, to collections (*e.g*. NEON Biorepository ^2^), to sequence repositories (*e.g*. GenBank ^3^). The practice of synthesizing data across multiple studies (*e.g*. exploring patterns such as taxonomic structuring, geographic trends, biotic and abiotic drivers and trends) relies on a consistency of data reporting, in terms of measurement units, specific IDs, nomenclatures, and well-specified terminology. For example, the use of trait data is widespread in ecological research, underpinning much of foundational exploration and approaches in ecological and evolutionary mechanisms. Thus, initiatives to standardize the wide variety of available ecological trait data exist, *e.g*. Schneider et al’s 2019 proposed Ecological Trait-data standard (ETS) ^4^.

The ability to synthesize and reuse data is particularly important for vector-borne disease (VBD) research. The risk of VBDs in people, livestock, wildlife, crops, and plants is currently increasing, in particular due to interactions with global change drivers ^5–8^. Understanding the shape and pattern of that risk, and potential additional risk for VBDs requires data on the underlying biological mechanisms of transmission. However, amassing the appropriate data to synthesize and analyze these essential model building blocks can be stymied by the sheer range of terminology, reporting styles, outputs, and simply a lack of a coherent framing to store them. While multiple databases for vector ecology data of many kinds exist, their scopes vary, as do their accessibility, and thus capacity for reuse and synthesis ^9^. Traits of arthropod vectors – measurable biological aspects of life-history, behavior, and vector competence – are integral to disentangling the complex mechanisms that underlie VBD transmission ^10^. Linking vector traits to transmission dynamics, in turn, is a crucial step in constructing useful mathematical frameworks and mechanistic models to predict disease dynamics and risk ^11–13^. While mechanistic models are undeniably powerful tools in the estimation of disease risk, the challenges of building and parameterising such models are also widely acknowledged. Chief among these is the sheer amount of data needed to parameterize models in biologically meaningful ways. The empirical data needed to derive realistic parameter estimates are typically collected through extensive experimentation in controlled laboratory settings. Thus, obtaining useful vector trait data, such as measurements of vector competence, fecundity, longevity, etc. across abiotic gradients (*e.g*., temperatures), is both financially and logistically costly to obtain. There is a clear benefit to leveraging large datasets synthesizing information from many sources (e.g. ^12,14–16^), yet the lack of a minimum information standard for reporting data generated by vector trait experiments impedes our capacity for aggregating data across collection efforts.

To ensure usability by the broader scientific community, datasets should adhere to FAIR Principles – Findable, Accessible, Interoperable, and Reusable – which are key components of good data management practices ^17^. Generally, the information shared will comprise two components, i) data, or measured traits and outcomes generated by experiments, and ii) metadata, or information about the origin of the data.

Here we present MIReVTD (Minimum Information standard for Reporting Vector Trait Data), a minimum information standard developed to accommodate vector trait experiment information in a flexible, transparent, and well documented database backbone, with accompanying metadata to facilitate data sharing and usability. Minimum information standards define a checklist of information minimally required to understand and reuse a biological dataset. They do not prescriptively define a specific set of field names or data types ^18^, but it is useful to provide examples in practice (data standards) which do, as illustration. Examples of minimum information standards include MIAPPE (Minimum Information About a Plant Phenotyping Experiment) ^19^; MIReAD (Minimum Information standard for Reporting arthropod Abundance Data) ^20^; and Wu et al’s minimum data standard for vector competence experiments ^21^. The minimum information standard we report here arose from efforts comprising two long-term research projects, one of which sought to define “what is a trait?” ^10^ for disease vectors, and the other is part of a long-term informatics project, VectorByte (VectorByte.org), that is building a database (VecTraits ^22^, doi.org/10.7274/28020782) containing the answer. The VecTraits database and format is an exemplar implementation and operationalization of the minimal information standard presented here, accommodating the minimum information needed, while providing flexibility for expanding fields to iterate across multiple axes of variation ^22^.

## Results

Among vector trait experiments and observations, there is considerable variation in vector trait data generated by independent studies, including which traits are measured, and the conditions under which they are measured. Due to the inherent complexity of data generated by vector trait experiments, this is not intended as a template for data collection, but rather a guide to what minimum information must be included when reporting outcomes, to ensure secondary use of data. At the most basic level, the minimum descriptor set for vector traits to maximize usability across studies are as follows:

### Organism

The genus and species of vector being studied and, if known, particular subspecies or lab strain. For individually measured data, this may also include some unique identifier to designate each individual or replicate (for example when multiple traits or timepoints are measured on the same individual organism). In transmission experiments, species or strain of pathogen must also be reported. Sex and life stage/age of the organism should be included.

### Trait Description

The vector trait being studied, how it was measured, the units of measurement used, and the frequency of observations. Ideally data should be in the least aggregated form available (*e.g*., measurements on individuals instead of means across individuals). When only means (or other summaries) are available, metrics of variability (*e.g*., standard error) and their descriptions (including sample sizes) should be included.

### Axes of Variation

Specify which abiotic (*e.g*., temperature) or biotic (*e.g*., food source) gradients were incorporated into the study, the frequency at which observations were made, and the units by which this variation was recorded. Multiple such covariates, many of which are biological “stressors” may be incorporated in each study. Further, any additional experimental settings (*e.g*., ambient temperature when temperature is not manipulated) should also be recorded. The sampling design should also be specified. For example, were trait measurements taken on multiple individuals at a single point in time, or were individuals tracked through time and measured across a gradient (several distinct treatments) such as time or temperature? Note that well annotated granular data makes experimental design self-evident.

The 3-component minimum information we outline here is expanded upon in Table 1 and in Box 1. Table 1 gives examples of data fields and the types of details, while Box 1 provides some more general suggestions and guidelines on formatting.

**Table 1.** Example data fields to capture minimum descriptors for vector trait experiments.

| Descriptor | Field(s) | Details | Recommendations | Examples |
| --- | --- | --- | --- | --- |
| **Organism** | Vector taxonomy | Genus and species of vector being studied, and if available, subspecies or laboratory strain | Be as specific as possible<br>Do not use abbreviations<br>If known, include lab or colony strain name<br>If relevant, a lab strain identification name / number / barcode | “*Aedes aegypti*”<br>“*Culex quinquefasciatus* Sebring colony”<br>“*Delphacodes kuscheli*”<br>“NR-44077 *Rhodnius prolixus*, Strain CDC” |
|  | Unique identifier | Designation of individuals or replicates in experiments, if applicable | Internal naming convention for study<br>Can be used to link related observations from a series or lab experiment<br>For example, a trait may be measured, on the same animal, at multiple ages - linked together by a unique (animal) identifier | “Mosquito1”<br>“tick_45” |
|  | Pathogen taxonomy<br>(when relevant, e.g. transmission studies) | Genus, species, and strain (if known) of pathogen | Be as specific as possible<br>Do not use abbreviations<br>If known, include viral or pathogen strain<br>If relevant, a lab strain identification name / number | “*Plasmodium falciparum* (N54 strain)”<br>“West Nile virus”<br>“Dengue virus DENV-4 (strain H241)”<br>“MRA-578 *Plasmodium falciparum*, Strain D10 PfM3' [D10-PfM3' (wt MSP-1 replacement)]” |
|  | Age / life stage | The age / life stage that was assayed | Be as specific as possible<br>Do not use abbreviations | “L1 larvae”<br>“3-4 day old post eclosion adults”<br>“nymphs” |
|  | Sex | The sex of the organism that was assayed |  | “males”<br>“unknown” |
|  |  |  |  | "mixed" |
| **Trait Description** | Vector trait | A detailed description of the trait being studied | Be as specific as possible, if appropriate specify life stage to avoid confusion<br>Avoid abbreviations | "mortality"<br>"lifespan"<br>"Fecundity"<br>"Development time - hatch to adult" |
|  | Value | A numerical measurement of the trait being studied | Provide units in separate field, or in column heading | 45<br>17<br>0.25 |
|  | Units of measurement | The units of measurement used to record values for trait data |  | "days"<br>"eggs laid"<br>"percent mortality"<br>"LT50" |
|  | Study location | Geographical location where field study was conducted, or where samples were collected | This should be recorded only for experiments where location could influence results, for example field studies or source location of collected individuals<br>E.g., locations where lab tests were run are not part of the experimental design, and may inadvertently suggest that local specimen strains were used<br>Be as detailed as possible, reporting latitude and longitude if available |  |
| **Axes of Variation** | Example field(s): | The abiotic experimental factor (variables) at which trait observations are measured | Common gradients include:<br>Temperature<br>Age<br>Location<br>Date | Separate different axes of variation into different columns |
|  | Units of measurement | The units of measurement for experimental factor |  | "Degrees celsius"<br>"Age, days post emergence" |

#### Box 1.

**General suggestions and guidelines for reporting traits data**

- Do not use abbreviations in data fields and especially not in field names - they introduce uncertainty.
- Avoid the use of more than one natural reporting language (e.g., mixing English and Standard Mandarin) as this can result in interpretation errors.
- Use numeric dates, preferably ISO 8601 format (*e.g*. YYYY-MM-DD)
- Provide data that is machine readable. For example, numerical data should be separated from its units into different fields. For more information on suggestions for reporting units, see Hanisch et al. ^23^
- Avoid diacritics (accent marks) and other special characters as they often lead to problems in reuse as some systems will not handle them correctly due to encoding differences.
- When providing the geographic location of data collection (for example when reporting a trait value across a geographic range), be as detailed as possible. Latitude and longitude are preferred, if available. Note that when lab tests were run and origin is not specified or is not part of the experimental design, this may inadvertently suggest that local specimen strains were used.
- The less processed the data are, the more reusable the data will be. We advocate providing per-organism data, when possible. For example, report data on the lifespan of each animal, instead of “average lifespan” or report each measure of a multiple-measure experimental design, linked with an appropriate unique (*e.g*. animal) identifier (e.g., number of eggs laid on a particular date by an individual mosquito).
- Save files in a text-based, non-proprietary format, preferably as a .csv file.
- **In Summary:** if a dataset is well reported and formatted, with all the minimal information, a secondary user should be able to understand from the data what experiment was performed just from looking at the data, including what organism was assayed, what was measured, under what condition(s), and what was the axis of variation.

The above outlines the bare minimum information needed to ensure that datasets are coherent and reusable beyond the original study. However, vector trait data may quickly become complex, necessitating additional detail and clarification to maintain their utility. In these instances, specificity matters. For example, when measuring wing length, along which axis is the measurement taken? When recording body mass, is it dry or wet mass, is weight taken for an intact insect, or is it wingless/legless mass, and for a single individual or averaged across multiple individuals? In addition to the basic required data fields, additional Trait Description fields should be added as needed to capture the dimensionality of information generated in a study. Further, there may be additional traits recorded in a study, which may influence the first trait described. In all of these instances, we can iterate the basic data inputs of Trait Description and Axis of Variation.

Metadata are equally vital for maintaining usability and interoperability of primary vector trait data. At minimum, reported metadata should include a full citation of the data source (*e.g*., this is typically a published paper), the name and contact for the person uploading the dataset to an online repository, and if relevant, the date on which any embargo on the dataset is lifted. We also advocate reporting data in the least-processed, most raw form. Generally, this means that for every experimental measurement or observation - that data point is represented individually in the reported data - and not as averages or derived values. As examples: reporting the lifespan of each individual animal in a mortality experiment instead of a LT50 or mean longevity.

### The importance of disaggregated data: an illustration

As an illustration of the importance of reporting individual level rather than group level information from vector trait experiments for mechanistic model parameterization, Figure 1 shows the difference in estimates generated for predicted juvenile development rate as a function of temperature, using group averages versus individual level data from Huxley et al’s ^16^ study on *Aedes aegypti*. Both datasets here are fitted with the same parametric function for the thermal performance curve (TPC), using the same Bayesian fitting algorithm and the same low information priors. Note how both the peak rate and the temperature bound estimates are impacted when averages are used in place of the original individual data, and the difference in errors around those estimates. In this case, strongly informative priors for the TPC model parameters (i.e., *T*_min_, *T*_max_) would need to be set when fitting to the averages in order to obtain fits that are comparable to the fit obtained for the individual level data. That is, extra outside information would need to be included to compensate for the loss of information that occurred when the averages were taken.

**Fig. 1.**
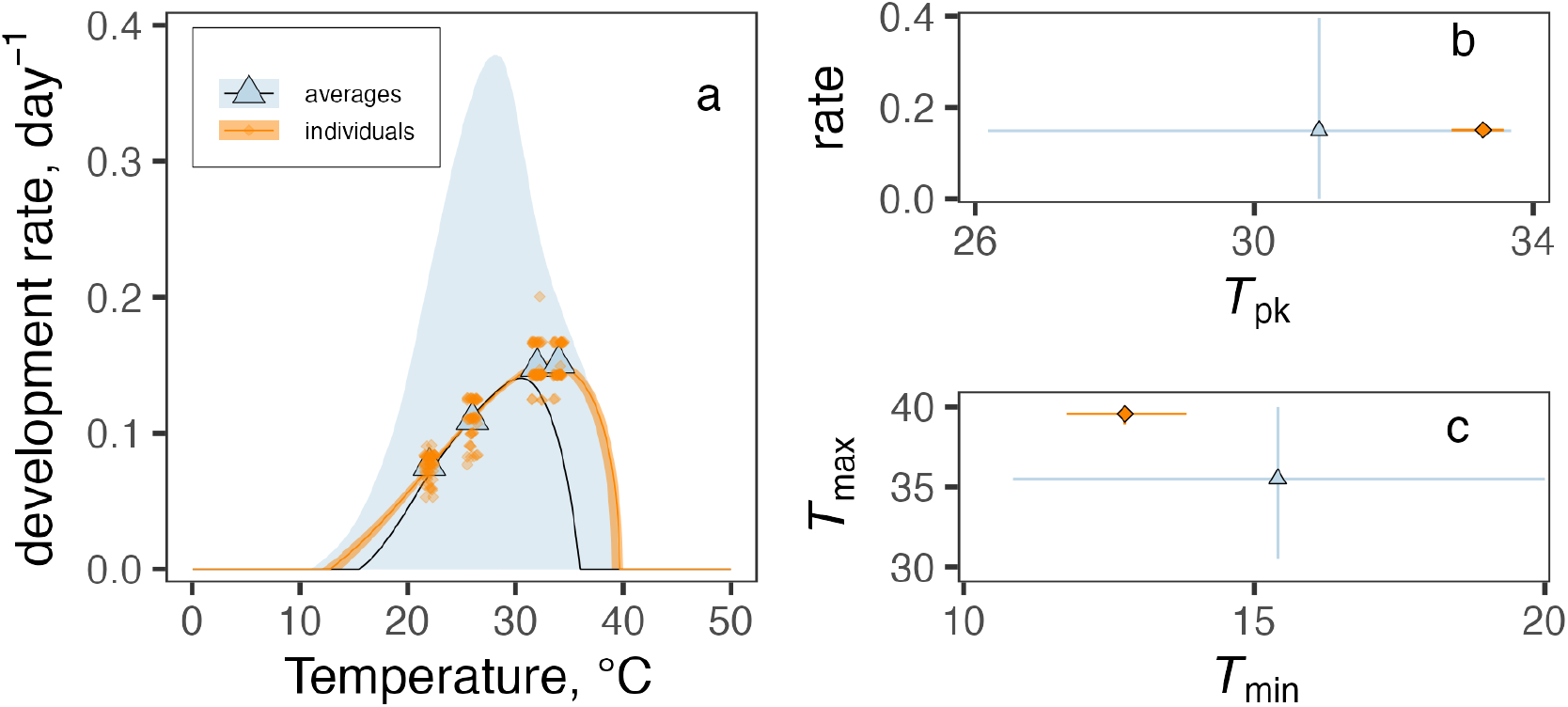
a–c. a. Example of some differences that can arise when TPCs (thermal performance curves) are fitted to single point averages versus individual-level observations when priors are set to be weakly informative (both fits use the same priors for all parameters). The trait fitted to here is juvenile development rate (inverse of duration from hatching to adult eclosion). Blue triangles denote averages; the blue bounds are the 95% credible intervals for the fitted central response (median). Orange diamonds are individual observations (slightly jittered); the orange bounds are the 95% credible intervals for the fitted central response (median). b. Differences between predicted development rate at its *T*_pk_ (i.e., the temperature at which a trait reaches its highest value) for TPCs fitted to averages (triangles) and individual observations (diamonds). c. Differences between predicted *T*_min_ and *T*_max_ for TPCs fitted to averages (triangles) and individual observations (diamonds). Points in b and c are median posterior estimates. Error bars in b and c are 95% highest posterior density (HPD) intervals for each parameter. TPCs were fitted to data from Huxley et al. 2022^16^ using the bayesTPC package ^24^.

### VecTraits Database

The VectorByte initiative (https://www.vectorbyte.org/) has worked to establish a global and openly accessible data hub to support vector research, which itself was inspired by earlier work on thermal traits from the BioTraits project ^25^. An outcome of VectorByte was the release of the VecTraits Database, an online platform for open hosting and sharing of biological vector trait data. Here, we use VecTraits to demonstrate an implementation of the minimum data standard, and additional suggested metadata capture. For data input, certain fields are required, by design, in part to maintain interoperability with other, earlier trait databases (especially BioTraits^26^, which focused on thermal traits). Fields are also required to satisfy the need for minimum information for reuse. Beyond the first set of required entry fields, iterations of fields for minimum standards (*e.g*., OrganismID, Trait Description) can be entered into VecTraits as needed, labeled as ‘interactors’. For example, vector competence and transmission studies should also include species or strain of pathogen used in the study, and this information would be recorded in VecTraits through the “interactor2” fields where appropriate. The interactor2 field is not required for uploading datasets into VecTraits because this may not apply to all experiments, but for studies that include pathogens, the minimum information standard described here indicates that this information is required to be reported. VecTraits thus tries to strike a balance between requiring sufficient fields be present and correctly inputted, and the flexibility to expand necessary columns of input to accommodate multiple axes of variations that may be included in a study or set of experiments. The current list of VecTraits field names and column definitions is spelled out, including examples of the data one would enter, the data format (TEXT, INTEGER, BOOLEAN, *etc*.), and restrictions on format (e.g. Not null, length <= 255 characters), at https://vectorbyte.crc.nd.edu/vectraits-columndefs.

### Example Dataset

Here we present an exemplar dataset retrieved from the VecTraits database to illustrate how this minimum information standard may be applied in practice (Fig 1). These data originated from a study by Huxley et al. 2022 ^27^ on the effects of larval competition and resource depletion on the temperature dependence of maximal population growth rates in the *Aedes aegypti* mosquito. This example demonstrates how the VecTraits database, with its own naming conventions and data entry fields, still complies with the minimum data standard while maintaining the flexibility needed to host data generated through complex experimental designs. By avoiding the use of a fixed template for all data columns, there is enough adaptability in the data entry process to expand columns as needed to capture requisite aspects of a given study that do not unilaterally apply to every experiment on vector traits. In this example, lifespan was recorded across a temperature gradient, which is a reportable “Axis of Variation” under the minimum data standard. This study also recorded juvenile lifespan (the duration from hatch to death or adult eclosion for all individuals in each sample population) at four initial resource concentration levels, which represents an additional “Axis of Variation” that is reportable under the minimum data standard, but is not universally applicable to most studies. In addition to meeting minimal data requirements, this example also highlights the collection of adequate metadata, where information on the published study where data originated is provided, as well as the name of the user who submitted the dataset to the VecTraits database. Note that while other data were collected in this study, including development time, longevity, and survival, these were entered as unique datasets with bespoke columns to reflect the dimensionality of the traits being measured, though these datasets are still linked through common metadata fields (*i.e*., citation and DOI).

**Fig. 2.**
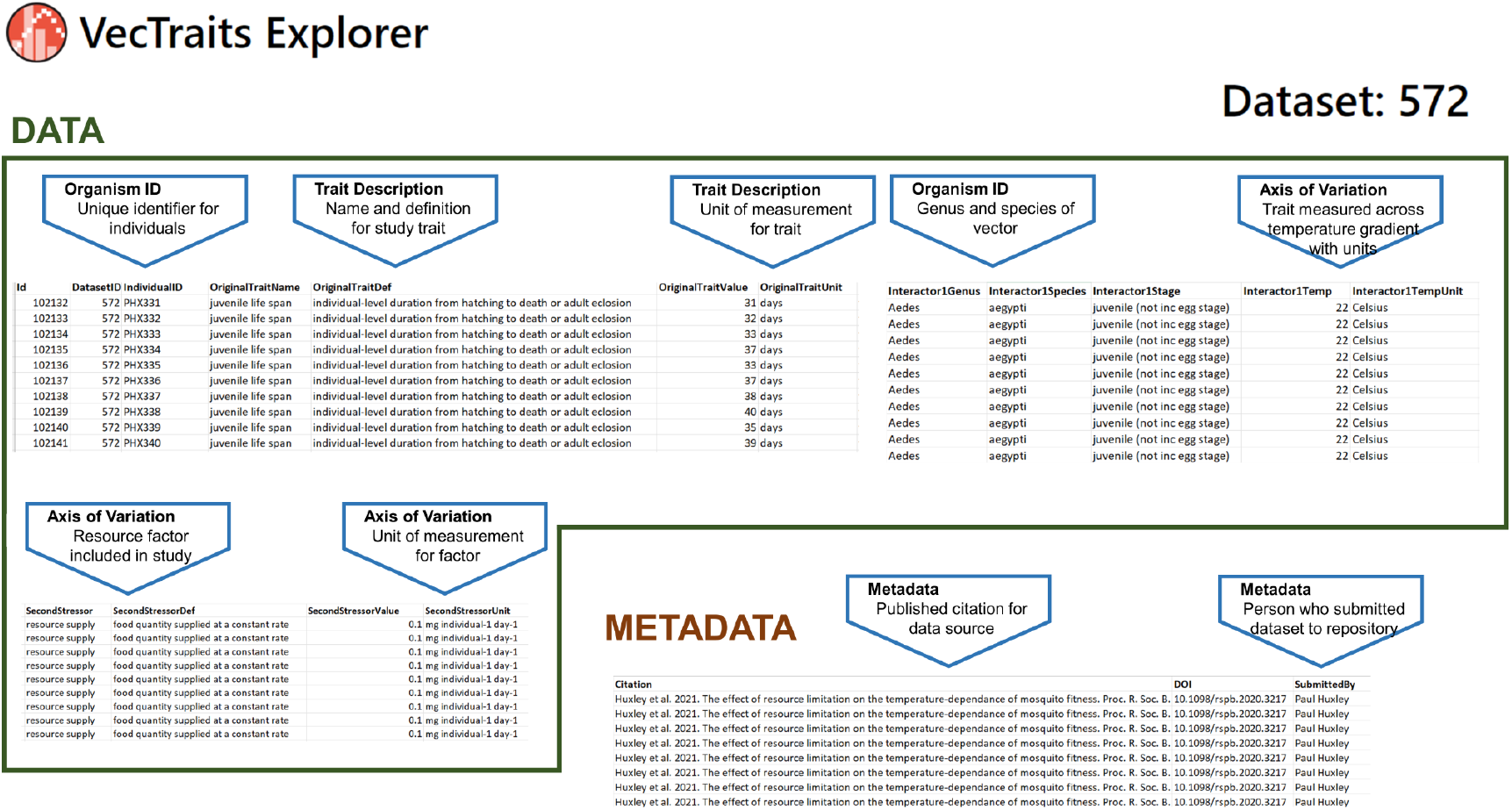
Example of data in VecTraits for a study measuring juvenile lifespan of *Aedes aegypti*. The trait in this dataset is juvenile life span and it is specified that this refers to surviving until death or adult eclosion. Each row contains a trait measure, number of days alive, for each individual mosquito in the experiment. This study included two stressors (temperature and resource level) and these are indicated as axes of variation. Measures and units are specified separately. The full citation is included to support source attribution.

## Discussion

Establishing minimum information standards for reporting and sharing vector traits data is an important step forward for maintaining the ‘reusability’ and FAIR-ness of experimental data. By emphasizing which elements of a study must be reported to ensure that datasets are usable beyond the original study, as opposed to providing a set template for data collection, the minimal information standard has the necessary flexibility to work with the multitude of experimental designs used to capture trait data, which are incredibly varied in purpose and format. Standardizing the reportable components of shared datasets will benefit the broader community of vector-borne disease researchers, particularly those whose work relies on experimentally derived data to parameterize models. The incorporation of variable trait data into modeling frameworks can deepen our understanding of transmission dynamics and expand the capacity for accurate disease modeling. Yet data-hungry methodological efforts are too often hindered by a lack of empirical data, which can result in unrealistic model predictions. Laboratory experiments designed to accurately measure vector traits across various Axes of Variation can be logistically demanding and resource intensive, effectively capping the sample size that is obtained from any single experiment. The need for empirical data to support VBD research has not gone unnoticed, and in recent years there have been great advances in the collection of large vector traits datasets, owing to government initiatives, innovations in empirical data collection, and the development of open data repositories. The increasing capacity to collect empirical data, and pool those observations across studies, underscores the pressing need for a cohesive set of minimum information data standards to facilitate secondary data analysis and promote FAIR Principles ^17^ in data sharing.

## Acknowledgements

Several authors were supported by CIBR: VectorByte: A Global Informatics Platform for studying the Ecology of Vector-Borne Diseases (SJR and CAL by NSF-DBI 2016265, LRJ and PH by NSF-DBI 2016264 and SSCR by NSF-DBI 2016282).

